# Profiles of cooperative brains: A discriminant analysis of cleaner and client fish monoaminergic responses to different social contexts

**DOI:** 10.1101/326843

**Authors:** Caio Maximino, Ana Cristina R. Gomes, Murilo S. de Abreu, Sónia C. Cardoso, Monica Lima-Maximino, Svante Winberg, Marta C. Soares

## Abstract

Vertebrate cognitive function requires a dynamic coordination of multiple specialized areas of the brain. The challenge here is to understand how these brain areas respond in dependence to the neurophysiological mechanisms in place, as to enable the successful processing of information. For instance, social and cooperative behaviour has been linked to the activation of some specific brain areas, mostly associated with reward processing. Here we evaluated a classic model system of cooperation between species of fish and compared datasets of brain monoaminergic response. We analysed by using multivariate discriminant analysis the exposure of cleaners, *Labroides dimidiatus,* to several social-related conditions, as well as the response of one client species, *Naso elegans,* to similar contexts. We demonstrate that the variable appraisal of each social challenge contributes to brain dopaminergic and serotonergic changes, in cleaners and clients, with both showing the diencephalon and optic tectum as main areas of metabolite response. The role of the serotoninergic system activation was mostly demonstrated at the diencephalon and cerebellum of cleaners, a response that was driven by mutualistic interaction, contact with client. Our current evidence is the first to jointly demonstrate the level of selective similarity in brain monoaminergic mechanisms that underlie fish mutualistic and social engagement, for both sides of these partnerships.

## 1. Introduction

The neural substrate of cooperative behaviour is gaining increasing attention to the recognition that cooperation is widespread in nature [Dugatkin 1997], especially on fish species [Soares et al. 2018]. Cleaner wrasses (*Labroides dimidiatus*) are mutualistic fish that engage in cooperative behaviour, inspecting the surface, gills and sometimes the mouth of visiting ‘client’ reef fish, eating ectoparasites, mucus, scales and dead or infected tissue [Bshary and Côté 2008; Soares 2017]. Cleaners display flexibility in adjusting their cleaning services in accordance with clients, varying between “cooperating” (removing ectoparasites) or “defecting” (eating fish mucus) [Bshary and Côté 2008].

There is limited evidence for a role of monoamines on cooperative behaviour. In healthy humans, tryptophan depletion reduced the level of cooperation shown by participants in the first day of an iterated Prisoner’s dilemma game [Wood et al. 2006]. In cleaner wrasses, fluoxetine and the serotonin (5-HT) 1A receptor agonist 8-OH-DPAT increased cooperative behaviour (raising engage in cleaning behaviour and the frequency of tactile stimulation), while blocking serotonin synthesis (by pCPA or antagonizing the 5-HT_1A_ receptor (WAY 100,635)) decreased cleaner’s cheating levels [Paula et al. 2015]. The blocking dopamine (DA) D1 receptors also increases cooperative behaviour (raising tactile stimulation and induces cleaners to initiate more interactions with clients) [Messias et al. 2016]. These effects appear to be mediated by the degree of familiarity with the client, with D1 antagonists increasing tactile stimulation only when interacting with novel clients [Soares et al. 2017]. It has been suggested [Soares et al. 2018] that DA modules predicted outcomes, with reductions in DA signalling contributing to an omission of reward (when the outcome is worse than predicted, e.g., lower probability of getting food during an interaction, or higher likelihood of being punished), suggesting that 5-HT and DA have opponent roles in tracking outcomes.

A relevant question focussing on the role of monoamines in this mutualistic system regards the importance of physical contact. Monoaminergic signalling has been proposed to mediate social behaviour in fish, increasing dopamine levels after visual contact with conspecifics [Saif et al. 2013]. In addition, in cleaning interactions, physical contact (known as massages) occurs frequently as it is provided by cleaners, and functions to benefit clients in terms of stress reduction [Soares et al. 2011]. Thus, the monoamines could act at various points of the cooperative and mutualistic interactions (e.g., client evaluation, motivation to approach the client, maintenance of cleaning engagement after initial contact, and during inspection [Soares 2017]). Moreover, DA is associated with the social behaviour network as to reinforce responses to salient social stimuli [O’Connell and Hofmann 2011]. Here we follow on recent published work of [Abreu et al. 2018b] and use this dataset to infer, only this time, by using multivariate discriminant analysis, on monoamine mediation on the motivation to approach and engage in social and mutualistic interactions by cleaners *L. dimidiatus* when introduced to a client or to another conspecific (in visual or physical access). Furthermore, we also used the previous published dataset of [Abreu et al. 2018a] to associate monoamines with the clients (*Naso elegans*) drive to being cleaned (when introduced to a cleaner) or to engage in social interactions (when introduced to another conspecific). We thus examined the level of selective similarity in brain monoaminergic mechanisms, by measuring brain monoamines DA and 5-HT and metabolites DOPAC (3,4-dihydroxyphenylacetic acid) and 5-HIAA (5-hydroxy indole acetic acid), at various brain areas (forebrain (FB - which included the olfactory bulbs and the telencephalon), diencephalon (DL), optic tectum (OT), cerebellum (CB) and brain stem (BS)), that may underlie fish mutualistic and social engagement, for both sides of these partnerships.

## Material and Methods

### 2.1. Experimental conditions

We analysed data from experiments conducted at the fish housing facilities of the Oceanário de Lisboa (Lisbon, Portugal). In these experiments were used 53 Indo-Pacific bluestreak cleaner wrasses, *Labroides dimidiatus* and 18 adults blond naso tang *Naso elegans* (family Acanthuridae; referred to as “clients”). For more information about the experimental conditions see [Abreu et al. 2018b; Abreu et al. 2018a]. The quantification of monoamines in fish occurred by high performance liquid chromatography with electrochemical detection (HPLC-EC). In brains were analyzed the DA and 5-HT, as well as the metabolites DOPAC (3,4-dihydroxyphenylacetic acid) and the 5-HIAA (5-hydroxy indole acetic acid), as described by [Øverli et al. 1999]. The experiments were carried out in accordance to the approved guidelines by the Oceanário de Lisboa and the Portuguese Veterinary Office (Direccao Geral de Veterinaria, license # 0420/000/000/2009).

#### 2.1.1. Cleaners perspective

Here, we used data from [Abreu et al. 2018b]. Cleaners were exposed to five different contextual treatments. Indeed, on each experimental day, one of the five following treatments was randomly allocated to each subject cleaner wrasse aquarium: a) conspecific (*L. dimidiatus*, n = 10), b) client (*N. elegans*, n = 11), c) conspecific inside another smaller aquarium (n = 12), d) client inside another smaller aquarium (n = 10) and e) ball (white ball, about 5 cm in diameter, which stayed at the bottom, completely sessile, n = 10), see Fig 1. Cleaners were exposed to treatments for the next 60 minutes. At the end of experiments, each subject cleaner wrasse was captured and immediately sacrificed with an overdose of tricaine solution (MS222, Pharmaq; 500– 1000 mg/L) and the spinal cord was sectioned. The brain was immediately dissected under a stereoscope (Zeiss; Stemi 2000) into five macro-areas: forebrain (FB - which included the olfactory bulbs and the telencephalon), diencephalon (DL), optic tectum (OT), cerebellum (CB) and brain stem (BS) (for a drawing of the major brain areas, please see [Abreu et al. 2018b; Abreu et al. 2018a]. Major brain areas were frozen with dry ice and the stored at - 80°C.

**Figure 1.**
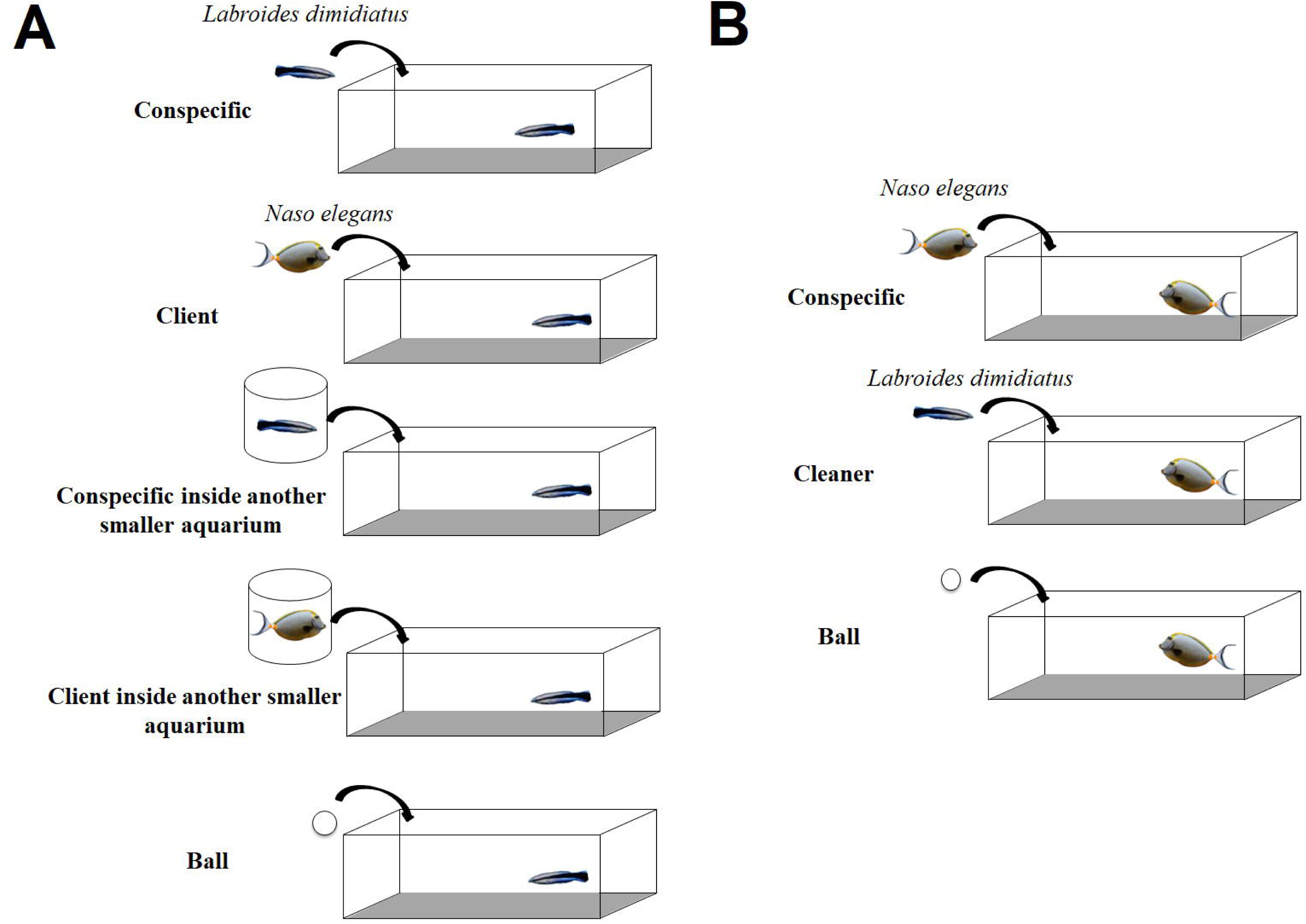
Experimental schematics. (A) cleaner and (B) client perspective.

#### 2.1.2. Clients perspective

We used data from [Abreu et al. 2018a], which included two of the treatments, and added a third context (ball group), similarly to use in cleaner perspective (see above). Three different contextual treatments were allocated to our subject clients (*N. elegans*): a) conspecific (*N. elegans*, n =7), b) a cleaner (*L. dimidiatus*, n = 7) and c) ball (same as above, n = 4) see Fig 1. Cleaners were exposed to treatments for the next 60 minutes. At the end of experiments, each subject client was captured and immediately sacrificed and brain micro-dissected as explained above.

### 2.2. Statistical analysis

We used multivariate discriminant analysis (MDA) to test for differences in the pattern of metabolite expression among the 5 social contexts for cleaners perspective, or the 3 social contexts for clients perspective. One set of MDA uses as data the concentrations of each monoamine (DA and 5-HT) and metabolite (DOPAC and 5-HIAA) at each brain area. This allows us to find the combination of metabolites and brain areas that discriminate the most the different social contexts. Another set of MDA uses as data the ratios of DOPAC/DA and 5-HT/5-HIAA as these will enable us to interpret the relation between monoaminergic production and its subsequent metabolic use. We interpret the two principal discriminant functions (DF1 and DF2) of each MDA based on trait loadings higher than 0.40 [Hair et al. 2010]. We assessed the significance of the discrimination within the entire set of social contexts using Wilk’s Lambda, and we used ANOVAs with post-hoc Tukey HSD tests to find which pairs of social context differ significantly in DF1 or DF2. All values were standardized using script available [Coghlan 2013] and normality was checked through q-q plots. All analyses were run with R (version 3.3.3; R Core Team, 2017) package car (version 2.1-4; [Fox and Weisberg 2011] and MASS (version 7.3-47; [Venables and Ripley 2010]).

Open science practices

Data for this manuscript can be found at our GitHub repository. (https://github.com/lanecunifesspa/cleaners).

## 3. Results

### 3.1. Cleaners perspective

Overall, social contexts differed in the levels of metabolite expression (Wilk’s lambda = 0.03, p = 0.001; Fig.2A) and in metabolite ratios (Wilk’s lambda = 0.23, p = 0.02; Fig.2 B) across the brain tissues of cleaners. For metabolite levels, DF1 and DF2 explained 83.9% of the variation between all the contexts. DF1 differed significantly between most pairs of contexts, except for the comparison between client vs. conspecific inside another aquarium, which instead were found to differ in DF2 (Fig. 2A, Table 1). DF1 was characterized mostly by strong negative loadings of DOPAC at the brain stem, and of DA, 5-HIAA and 5-HT at the diencephalon of cleaners (loadings <-0.40; all other loadings < |0.29|; Fig. 2C and Table 2). DF2 was characterized mostly by a strong positive loading of DA at the optic tectum (loading = 0.58; all other loadings < |0.36|; Fig. 2C and Table 2).

**Table 1.** Pairwise correlations of contextual groups’ differences that are significantly explained by each of the discriminant functions, for cleaners. * significant relationships (p < 0.05)

|  | Monoamine |  | Monoamine ratio |  |
| --- | --- | --- | --- | --- |
|  | DF1 | DF2 | DF1 | DF2 |
| **Conspecific vs. Control** | 0.02* | 6.85E-06* | 1.00 | 0.75 |
| **Conspecific inside**<br>**aquarium vs. Control** | 5.07E-11* | 0.98 | 0.63 | 0.34 |
| **Heterospecific vs. Control** | 2.92E-07* | <0.01* | <0.01* | 0.42 |
| **Heterospecific inside**<br>**aquarium vs. Control** | 0.02* | 0.37 | 0.11 | 0.51 |
| **Conspecific inside**<br>**aquarium vs. Conspecific** | 2.47E-05* | 5.55E-07* | 0.58 | 0.99 |
| **Heterospecific vs.**<br>**Conspecific** | 0.03* | 0.65 | <0.01* | 0.05* |
| **Heterospecific inside**<br>**aquarium vs. Conspecific** | 1.48E-06* | <0.01* | 0.21 | 0.07 |
| **Heterospecific vs.**<br>**Conspecific inside**<br>**aquarium** | 0.16 | 2.23E-05* | 0.08 | <0.01* |
| **Heterospecific inside**<br>**aquarium vs. Conspecific**<br>**inside aquarium** | 2.49E-13* | 0.12 | <0.01* | 0.01 |
| **Heterospecific inside** | 7.95E-12* | 0.05 | 8.49E-07* | 1.00 |
| **aquarium vs.** |  |  |  |  |
| **Heterospecific** |  |  |  |  |

**Table 2.** Coefficients of each monoaminergic metabolites, synthesized at each brain area, for both clients and cleaners. DF1 and DF2 explained variation is, respectively, 87.4% and 12.6% for clients, and 63.8% and 20.1% for cleaners. * important metabolites (correlational coefficient > |0.40|).

|  | Cleaners |  | Clients |  |
| --- | --- | --- | --- | --- |
|  | DF1 | DF2 | DF1 | DF2 |
| **BS_DOPAC** | *-0.419 | 0.137 | -0.321 | 0.220 |
| **BS_5-HIAA** | 0.185 | -0.110 | -0.147 | 0.011 |
| **BS_DA** | 0.196 | 0.364 | -0.219 | 0.116 |
| **BS_5-HT** | 0.233 | 0.102 | 0.053 | 0.180 |
| **DL_DOPAC** | -0.087 | -0.257 | *0.427 | 0.062 |
| **DL_5-HIAA** | *-0.536 | 0.116 | *-0.401 | 0.085 |
| **DL_DA** | *-0.503 | 0.022 | -0.214 | *0.503 |
| **DL_5-HT** | *-0.588 | 0.167 | -0.154 | 0.396 |
| **FB_DOPAC** | -0.153 | -0.305 | -0.363 | -0.107 |
| **FB_5-HIAA** | 0.167 | 0.212 | -0.228 | -0.329 |
| **FB_DA** | -0.068 | -0.272 | -0.267 | -0.081 |
| **FB_5-HT** | -0.276 | 0.190 | 0.166 | -0.302 |
| **OT_DOPAC** | 0.045 | -0.260 | *-0.444 | 0.117 |
| **OT_5-HIAA** | 0.187 | -0.012 | -0.041 | 0.007 |
| **OT_DA** | -0.165 | *0.582 | -0.214 | 0.051 |
| **OT_5-HT** | 0.075 | 0.233 | 0.011 | -0.211 |
| **CB_DOPAC** | 0.067 | -0.316 | - | - |
| **CB_5-HIAA** | -0.201 | -0.235 | - | - |
| **CB_DA** | 0.050 | -0.151 | - | - |
| **CB_5-HT** | -0.292 | -0.183 | - | - |

**Figure 2.**
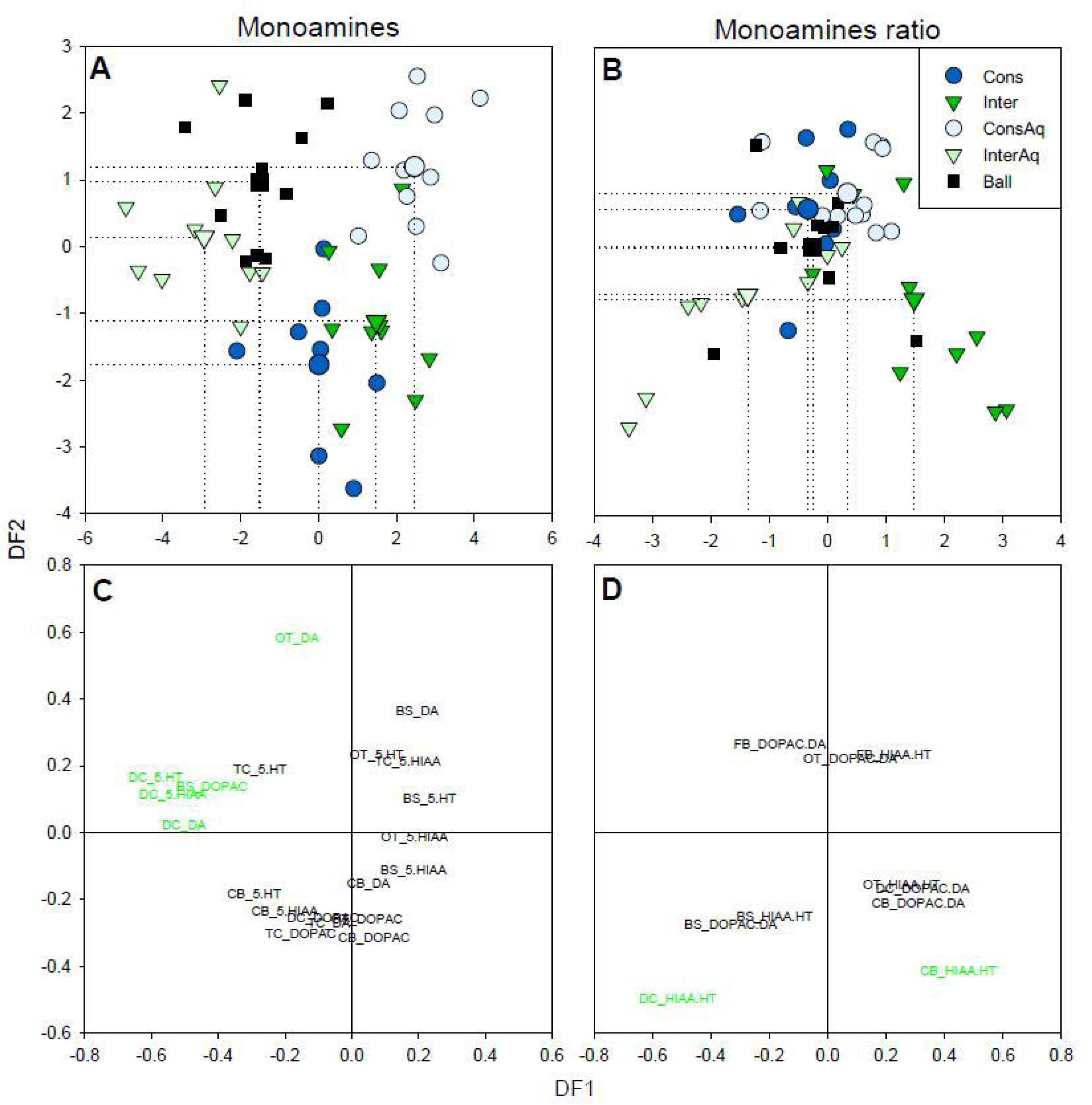
Cleaner’s discriminant analysis plot of DF1 versus DF2 to visualize how the discriminant functions discriminate between contextual treatments, for both monoaminergic metabolites and monoaminergic ratios, synthesized at each brain area. Bigger symbols with dotted drop line in the linear discriminant plot indicate centroid of each group. DF1 and DF2 explained variation is, respectively, for monoaminergic metabolites analysis (A) 63.8% and 20.1% and for monoaminergic ratios (B), 50.8% and 25.1%. (C) and (D), Trait loadings of each variable on DF1 and DF2; the most relevant metabolites to interpret DF1 or DF2 (trait loadings > 0.40) are shown in green. See the methods section for acronyms.

When using instead ratios of metabolite expression in the brain tissues of cleaners, DF1 and DF2 together explained 75% of the variation among contexts. DF1 differed significantly between cleaners that were prevented to interact with clients (inside a smaller aquarium) from those that could engage freely with clients (Fig 2B; Table 1). DF1 also differed between fully interactive contexts (i.e. interaction with clients) and either controls or interaction with conspecifics, and between clients in separate aquaria and conspecifics in separate aquaria (Fig 2B; Table 1). DF2 differed only between interaction with clients and either conspecifics in separate aquaria or conspecifics (Fig 2B; Table 1). DF1 was characterized mostly by loadings of 5-HIAA/5-HT ratios at the diencephalon and at the cerebellum, although in different directions (loadings = -0.51 and 0.45, respectively; all other loadings <|0.33|; Fig. 2D and Table 3). DF2 was characterized mostly by a strong negative loading of 5-HIAA/5-HT ratio at the diencephalon and cerebellum (loading = -0.50 and -0.41, respectively; all other loadings < |0.27|; Fig. 2D and Table 3).

**Table 3.** Coefficients of each monoaminergic ratios, synthesized at each brain area, for both clients and cleaners. DF1 and DF2 explained variation is, respectively, 70.2% and 29.8% for clients, and 50.8% and 25.1% for cleaners. * important metabolites (correlational coefficient > |0.40|).

|  | Cleaners |  | Clients |  |
| --- | --- | --- | --- | --- |
|  | DF1 | DF2 | DF1 | DF2 |
| **BS_DOPAC/DA** | -0.332 | -0.276 | -0.081 | -0.033 |
| **BS_5-HIAA/5-HT** | -0.182 | -0.252 | -0.341 | 0.310 |
| **DL_DOPAC/DA** | 0.324 | -0.166 | 0.710 | -0.269 |
| **DL_5-HIAA/5-HT** | *-0.514 | *-0.495 | -0.138 | -0.514 |
| **FB_DOPAC/DA** | -0.165 | 0.265 | 0.136 | -0.357 |
| **FB_5-HIAA/5-HT** | 0.227 | 0.233 | -0.551 | -0.012 |
| **OT_DOPAC/DA** | 0.077 | 0.220 | -0.278 | 0.380 |
| **OT_5-HIAA/5-HT** | 0.252 | -0.156 | 0.309 | 0.526 |
| **CB_DOPAC/DA** | 0.309 | -0.210 | - | - |
| **CB_5-HIAA/5-HT** | *0.446 | *-0.413 | - | - |

### 3.2. Clients perspective

Regarding clients, the exposure to different social contextual treatments was solely found to differ significantly in the levels of metabolite expression (Wilk’s lambda = 0.01, p = 0.02), showing a clear separation between the individuals exposed to different contexts (Fig. 3A) but not for metabolite ratios (Wilk’s lambda = 0.42, p = 0.51, Fig 3B).

**Figure 3.**
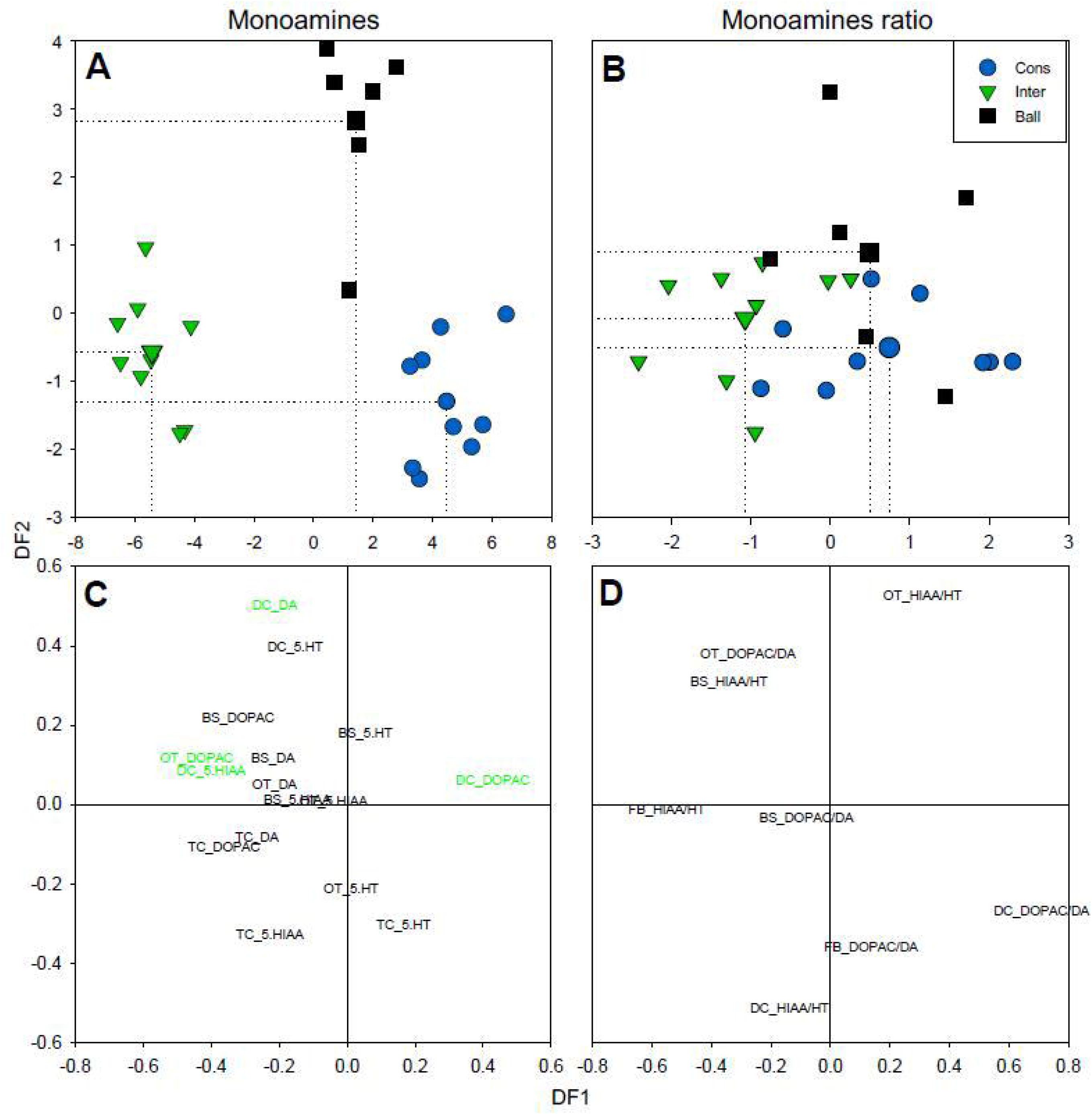
Clients’ discriminant analysis plot of DF1 versus DF2 to visualize how the discriminant functions discriminate between contextual treatments, for both monoaminergic metabolites and monoaminergic ratios, synthesized at each brain area. Bigger symbols with dotted drop line in the linear discriminant plot indicate centroid of each group. DF1 and DF2 explained variation is, respectively, for monoaminergic metabolites analysis (A), 87.4% and 12.6% and for monoaminergic ratios (B), 70.2% and 29.8%. In (C) and (D), relevant metabolites (correlational coefficient > 0.40) are shown in green. See the methods section for acronyms.

For monoaminergic metabolite production, DF1 and DF2 explained 99% of the variation and differ significantly among all pairs of contexts, except for the context of a cleaner with a client, which was not significant for DF2 (Table 4). DF1 seems to be explained by two areas of client fish *N. elegans*: the diencephalon (varying in terms of DOPAC and 5-HIAA, in opposite directions) and the optic tectum (solely for DOPAC; Fig. 3C, Table 2), while DF2 seems to be explained by the DA on the diencephalon (Fig. 3C, Table 2). Surprisingly, no differences were found in the overall ratio patterns (monoaminergic ratios DOPAC/DA and 5-HIAA/5-HT; Wilk’s lambda = 0.42, p = 0.51).

**Table 4.** Pairwise correlations of contextual groups’ differences that are significantly explained by each of the discriminant functions, for clients. * significant relationships (p < 0.05)

|  | Monoamine |  | Monoamine ratio<br>DF1 and DF2 |
| --- | --- | --- | --- |
|  | DF1 | DF2 |  |
| **Heterospecific vs. Control** | 4.67E-11 * | 6.26E-06* | - |
| **Conspecific vs. Control** | 2.96E-05* | 3.38E-07* | - |
| **Conspecific vs.**<br>**Heterospecific** | 4.72E-14* | 0.29 | - |

## 4. Discussion

Discriminant analysis is a powerful statistical tool that enabled us to determine the contribution of social and mutualistic behaviour to the modulation of brain monoaminergic levels of cleaners and clients, and specifically those linked to the activation of some localized brain areas. We found that brain monoaminergic response to social challenges shared interesting similarities for cleaners and clients, with most relevant alterations occurring at the diencephalon and optic tectum, in reference to DAergic and 5-HTergic change. While client fish clearly responded to differences posed by conspecifics compared to heterospecifics, we discovered that it was cleaners’ 5-HT system that mostly contributed to separate mutualistic activities from all others. This interaction is changed crucially whenever these individuals were given the possibility to physically engage in cleaning (occurring at the cerebellum) or were prevented to interact (occurring at the diencephalon) [Abreu et al. 2018b].

Broad analysis of cleaners’ monoaminergic response in respect to all different treatments suggests the distinction of 3 groups: a first with inaccessible clients and ball group, a second composed by inaccessible conspecifics and a third that included conspecifics and heterospecifics that were fully accessible. Specifically, the DF1 axis represented most of the variation observed, being mostly associated with DA, 5-HT and 5-HIAA at the diencephalon and DOPAC at the brain stem; while the DF2 mostly distinguished those stimuli that cleaners could physically access from those they could not, being these last group associated with DA levels at the optic tectum. It is usually considered that monoaminergic increase (DA or 5-HT) means an increment in production while the elevation of metabolite levels (DOPAC or 5-HIAA) signals for the actual system activation and monoamine usage (see [Saif et al. 2013]). One potential explanation for the individualized response of the DAergic system at the optic tectum, which corresponds with the ball group, is cleaners’ visual stimulation, in the absence of any other stimulation; while the DOPAC at the brain stem should be related to an increase of motor activity, as cleaners moved around the smaller client-aquaria. Moreover, the remaining relevant 5-HIAA varied at the diencephalon and all in response to the inability to interact with clients. While there was a clear distinction between the cleaners that could freely contact with stimuli and those that were unable (DA stimulation at the optic tectum), it was solely by looking at the metabolite/neurotransmitter ratios that the differences between being able to clean or not, were finally disclosed: the DF2 axis distinguishing between mutualistic and non-mutualistic activities and the DF1 discriminating between the actual physical contact, of those that were able to fully inspect a client (stronger association with 5-HIAA/5-HT at the cerebellum) and those that could not engage in cleaning (stronger association with 5-HIAA/5-HT at the diencephalon). These points for an extra turnover of 5-HTergic metabolites at the diencephalon whenever cleaners fail to achieve their needs and an increase of neurotransmission at the cerebellum during mutualistic activities.

Clients brain monoaminergic levels was also tested in respect to 3 main social challenges, revealing interesting similarities but also some relevant dissimilarities with cleaners’ overall response; for instance, all 3 groups clearly discriminated, with the DF1 axis again distinguishing the majority of the variation observed, being strongly associated with 5-HIAA at the diencephalon and DOPAC at the optic tectum but also with DOPAC at the diencephalon, in an opposite direction; while the DF2 mostly selecting living from non-living stimuli, being this distinction mostly related with DA levels at the diencephalon, in the presence of non-living stimuli. Thus, in the control treatment, clients seem to experience a general stimulation of the diencephalic DAergic system at the diencephalon contrasting with the overall stimulation at the optic tectum in cleaners.

Overall, teleost fish (and other vertebrate groups) cognitive function requires a dynamic coordination of multiple specialized areas of the brain. We demonstrate that the variable appraisal of each social challenge contributes to brain dopaminergic and serotonergic changes, in cleaners and clients, with both showing the diencephalon and optic tectum as main areas of monoaminergic response. However, the role of the serotoninergic system activation was mostly demonstrated at the diencephalon and cerebellum of cleaners, a response that was driven by the exposure to mutualistic stimuli. Our current results are the first to jointly demonstrate the level of selective similarity in brain monoaminergic mechanisms that underlie fish mutualistic and social engagement, for both sides of these partnerships.

## Author contributions

M.C.S. conceived the study. A.C.R.G. performed statistical analysis. All authors participated in writing and provided comments and criticism and gave final approval for publication. The authors declare no competing interests.

### Acknowledgements

We thank Gonçalo Cardoso for helpful discussions during the statistical analysis and comments on earlier drafts of this manuscript.

## References

Abreu MS, Messias JPM, Thörnqvist PO, Winberg S, Soares MC (2018a) Monoaminergic levels at the forebrain and diencephalon signal for the occurrence of mutualistic and conspecific engagement in client fish. Scientific Reports.

Abreu MS, Messias JPM, Thörnqvist PO, Winberg S, Soares MC (2018b) The variable monoaminergic outcomes of cleaner fish’ brains when facing different social and mutualistic contexts. PeerJ.

Bshary R, Côté I (2008) New perspectives on marine cleaning mutualism. In: Fish behaviour. Enfield: Science Publishers.

Coghlan A (2013) A little book of R for multivariate analysis. Wellcome Trust Sanger Institute, Cambridge, UK:1–47.

Dugatkin LA (1997) The evolution of cooperation. Bioscience 47:355–362.

Fox J, Weisberg S (2011) An R Companion to Applied Regression: SAGE Publications.

Hair JF, Black WC, Babin BJ, Anderson RE (2010) Applications of SEM. Multivariate data analysis. In: Pearson, Upper Saddle River.

Messias JPM, Paula JR, Grutter AS, Bshary R, Soares MC (2016) Dopamine disruption increases negotiation for cooperative interactions in a fish. Scientific Reports 6:20817.

O’Connell LA, Hofmann HA (2011) The Vertebrate mesolimbic reward system and social behavior network: A comparative synthesis. The Journal of Comparative Neurology 519:3599–3639.

Paula JR, Messias JP, Grutter AS, Bshary R, Soares MC (2015) The role of serotonin in the modulation of cooperative behavior. Behavioral Ecology 26:1005–1012.

Saif M, Chatterjee D, Buske C, Gerlai R (2013) Sight of conspecific images induces changes in neurochemistry in zebrafish. Behavioural Brain Research 243:294–299.

Soares MC, Oliveira RF, Ros AFH, Grutter AS, Bshary R (2011) Tactile stimulation lowers stress in fish. Nature Communications 2:534.

Soares MC, Santos TP, Messias JPM (2017) Dopamine disruption increases cleanerfish cooperative investment in novel client partners. Royal Society Open Science 4.

Soares MC (2017) The neurobiology of mutualistic behavior: the cleanerfish swims into the spotlight. Frontiers in behavioral neuroscience 11:191.

Soares MC, Cardoso SC, Carvalho TdS, Maximino C (2018) Using model fish to study the biological mechanisms of cooperative behaviour: A future for translational research concerning social anxiety disorders? Progress in Neuro-Psychopharmacology and Biological Psychiatry 82:205–215.

Venables WN, Ripley BD (2010) Modern Applied Statistics with S: Springer New York.

Wood RM, Rilling JK, Sanfey AG, Bhagwagar Z, Rogers RD (2006) Effects of Tryptophan Depletion on the Performance of an Iterated Prisoner’s Dilemma Game in Healthy Adults. Neuropsychopharmacology 31:1075.

Øverli Ø, Harris CA, Winberg S (1999) Short-Term Effects of Fights for Social Dominance and the Establishment of Dominant-Subordinate Relationships on Brain Monoamines and Cortisol in Rainbow Trout. Brain, Behavior and Evolution 54:263–275.

